# A comparison of clustering models for inference of T cell receptor antigen specificity

**DOI:** 10.1101/2023.08.04.551940

**Authors:** Dan Hudson, Alex Lubbock, Mark Basham, Hashem Koohy

## Abstract

The vast potential sequence diversity of TCRs and their ligands has presented an historic barrier to computational prediction of TCR epitope specificity, a holy grail of quantitative immunology. One common approach is to cluster sequences together, on the assumption that similar receptors bind similar epitopes. Here, we provide an independent evaluation of widely used clustering algorithms for TCR specificity inference, observing some variability in predictive performance between models, and marked differences in scalability. Despite these differences, we find that different algorithms produce clusters with high degrees of similarity for receptors recognising the same epitope. Our analysis highlights an unmet need for improvement of complex models over a simple Hamming distance comparator, and strengthens the case for use of clustering models in TCR specificity inference.

## Introduction

T lymphocytes recognise peptide epitopes presented at the cell surface by Major Histocompatibility Complexes (MHC) in jawed vertebrates [1]. Recognition is mediated by diverse heterodimeric *α* or *β* TCR domains positioned on the T cell surface. The chains of the more common *αβ* TCR contain variable (V), joining (J) gene segments, constant (C) regions, and an additional diversity (D) segment in the *β* polypeptide.

Each T cell expresses many copies of a single TCR, which bind to peptide-MHC (pMHC) via the complementarity determining regions (CDR) 1-3 of the TCR [2]. Productive TCR engagement triggers a context-dependent signalling cascade, which in turn promotes activation and differentiation of diverse immune effector cells [3].

The central role of the TCR in immune surveillance and response to disease has encouraged efforts to decode the rules of TCR-pMHC binding. Determination of specificity, or “receptor de-orphanisation”, can be achieved experimentally using sequencing and repertoire analysis, or with functional, multimer binding, or TCR screening methods, reviewed in [4], [5]. However, the ability to accurately predict the cognate epitope of any TCR *in silico* could vastly accelerate our understanding of fundamental and translational T cell biology [6].

The availability of large repositories of TCR sequences and their known ligands has enabled the development of two major families of computational model for prediction of TCR antigen specificity: Supervised Predictive Models (SPMs) and Unsupervised Clustering Models (UCMs) (**Fig. 1**)[6]. These families are representative of two distinct approaches to machine learning. In *supervised learning*, predictive models are trained on a set of input instances having a known label (in this case, the cognate epitope for a given TCR). In *unsupervised learning*, models learn the underlying statistical features or patterns of a dataset to differentiate between input TCRs, applying techniques such as clustering or dimensionality reduction.

**Figure 1.**
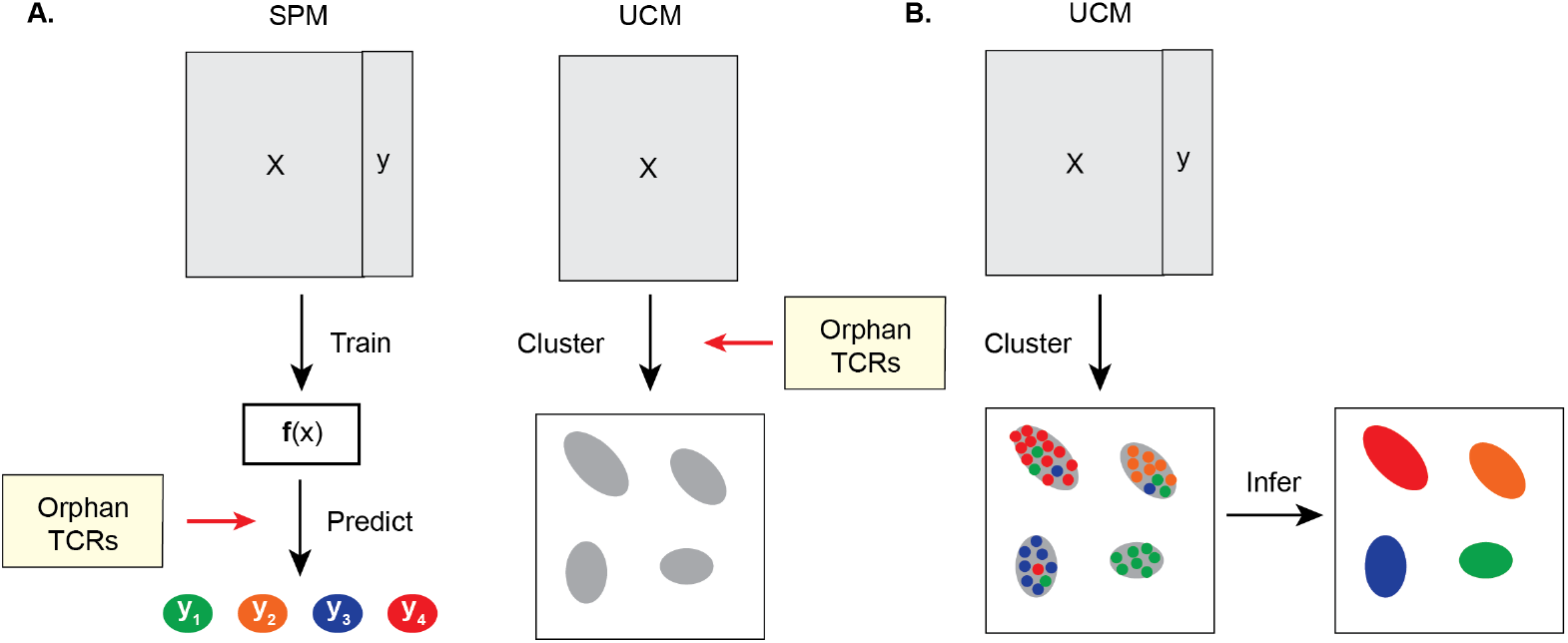
Supervised and unsupervised learning in T cell epitope specificity inference. A) SPMs (*left*) fit a predictive function **f** (x) to training data having an independent variable *X* (TCR sequences and other features) and dependent variable *y* (epitopes or pMHC complexes). This function may then be applied to predict the cognate epitopes of orphan TCRs. UCMs (*right*) generate a mapping from TCR sequences to a cluster allocation, such that each TCR is assigned to one or more clusters having common epitope specificity. B) When applied to datasets including full or partial epitope labels, UCMs may be used to predict TCR epitope specificity by assigning the most frequent epitope of a cluster as the predicted binder for all TCRs in that cluster.

The use of deep neural networks (DNNs) including large language models and convolutional neural networks has contributed significantly to recent improvements in UCM and SPM performance [7]–[11]. Despite these advances, no publicly available SPM is yet capable of accurately predicting the specificity of TCRs recognising “unseen” epitopes that were not encountered during model training [8], [12]. This is likely due at least in part to the limited volume of experimentally determined receptor-epitope pairs, which constitutes just a small fraction of the vast theoretical diversity of TCRs [6], [13].

Unlike SPMs, UCMs do not require receptor-ligand pairs as an input, but group similar TCRs together on the assumption that receptors having similar sequences will bind similar epitopes [14], [15]. UCMs can therefore be applied to identify clusters of similar TCRs irrespective of whether their cognate pMHC has been observed before.

This is of particular use in an era when bulk and single-cell sequencing experiments can yield thousands of unique TCRs per sample, by applying UCMs to shortlist TCRs of interest for later experimental de-orphanisation. Such approaches have been successfully applied to identify and characterise TCRs associated with mycobacterial and viral infection, cancer, and autoimmune disease [15]–[20].

UCMs take as their input single or paired TCR CDR3 nucleotide or amino acid sequences, with or without V and J gene usage information, and return a mapping of sequences to unique clusters. This has historically been achieved using some form of distance measure, typically either direct sequence similarity and/ or the frequency enrichment of short sequence snippets (*kmers*) compared to a reference dataset. Recent approaches leverage DNNs to generate a compressed numeric representation of the input TCR as a precursor to clustering [11], [21]–[23]. We dub such models DNN-UCMs to differentiate them from traditional distance-based UCMs such as GLIPH [15] and tcrdist [14], which were published simultaneously in 2017. A critical advantage of these “traditional” UCMs is that they do not require large volumes of training data, a hallmark and limitation of DNNs.

When some or all of the cognate epitopes of a given TCR dataset are known, UCMs can theoretically be used to infer epitope specificity (**Fig. 1B**). However, there has to date been no independent benchmarking study of UCMs as predictors of TCR specificity, despite their widespread use in field. In the present work, we compare the predictive performance of five commonly used UCMs on sets of known TCR-epitope pairs. We then extend our analysis to qualitative comparison of the clusters formed, practical considerations including runtime speed, and finally the impact on inference of introducing noise from synthetic background TCRs.

## Results

Benchmarking analyses were performed on paired *αβ* TCRs data drawn from VDJdb, a large, public, curated source of TCRs of known epitope specificity [24]. Model performance was analysed for data subsets generated by retaining epitopes having 10, 50, 100, 500, or 1000 cognate TCRs (datasets V10, V50, V100, V500, and V1000 respectively, **Table S1**). Instances were randomly down sampled after pre-processing, such that each experimental run was performed on the same number of TCR sequences per epitope, and all models were applied to *α* or *β* chain selections from the same set of paired TCRs. Sampling was repeated to account for sample variance between epitopes (**Table S2**). We present performance on *α* or *β* chain selections independently (see **Discussion** and **Limitations**).

Five open-source models were identified from the literature for which a python implementation was readily available: ClusTCR, GIANA, GLIPH2, iSMART, and tcrdist3 [17], [20], [25]–[27]. Three baseline models were added: A Hamming distance model that grouped together sequences having identical length and differing by not more than one amino acid; a CDR3 length-based model, and a random baseline. Details of model implementations are provided in **Methods**, and a summary of the respective methodologies in **Section S1**. As tcrdist3 generates a distance measure but does not explicitly cluster instances, a scikit-learn implementation of DBSCAN [28] was used to group distance matrices produced with tcrdist3, consistent with [25] and following comparison of model performance with different model implementations and clustering approaches (**Fig. S1**). Hamming, GLIPH2, tcrdist3, and iSMART implementations were adapted from the ClusTCR python package [25] and run using default parameters. The comparative analytical framework and model datasets are made freely available at *https://github.com/hudsondan/tcr-scapes*.

UCMs have historically been evaluated using cluster quality metrics such as purity, consistency, diversity, and retention (detailed in **Methods**). By applying each model to sets of TCRs having known specificity, we were able to combine these metrics with direct measures of predictive capacity including accuracy, precision, recall, and F1-score, following the schema depicted in **Fig. 1B**. Observing a positive correlation between performance according to cluster purity, consistency, adjusted mutual information, precision, recall, and F1-score (**Fig. S2**), we present results for F1-score alone (see **Supplementary Tables**), weighting F1-scores to account for model-specific class imbalance.

Differential model performance was first inspected at a global level for datasets V10 to V1000 combined, grouping over *α* and *β* chain selections and over all epitopes (**Fig. 2A-B** and **Table S3**). All study models outperformed length and random baselines (p*<*0.0001). However, whilst tcrdist3 generally performed well, absolute differences between many UCMs and a simple Hamming models were minimal (1-3% when accounting for 95% confidence intervals) (**Table S3**). Despite the apparent statistical significance of many comparisons between models (**Fig. 2B**), the relative performance of each model was sensitive to the TCR chain selection (**Fig. S3** and **Table S3**). Furthermore, rankings did not hold across epitopes (**Fig. 2C** and **Table S4**), nor when applied to a discrete set of 509 TCRs drawn from McPas-TCR (**Fig. S4** and **Table S5**).

**Figure 2.**
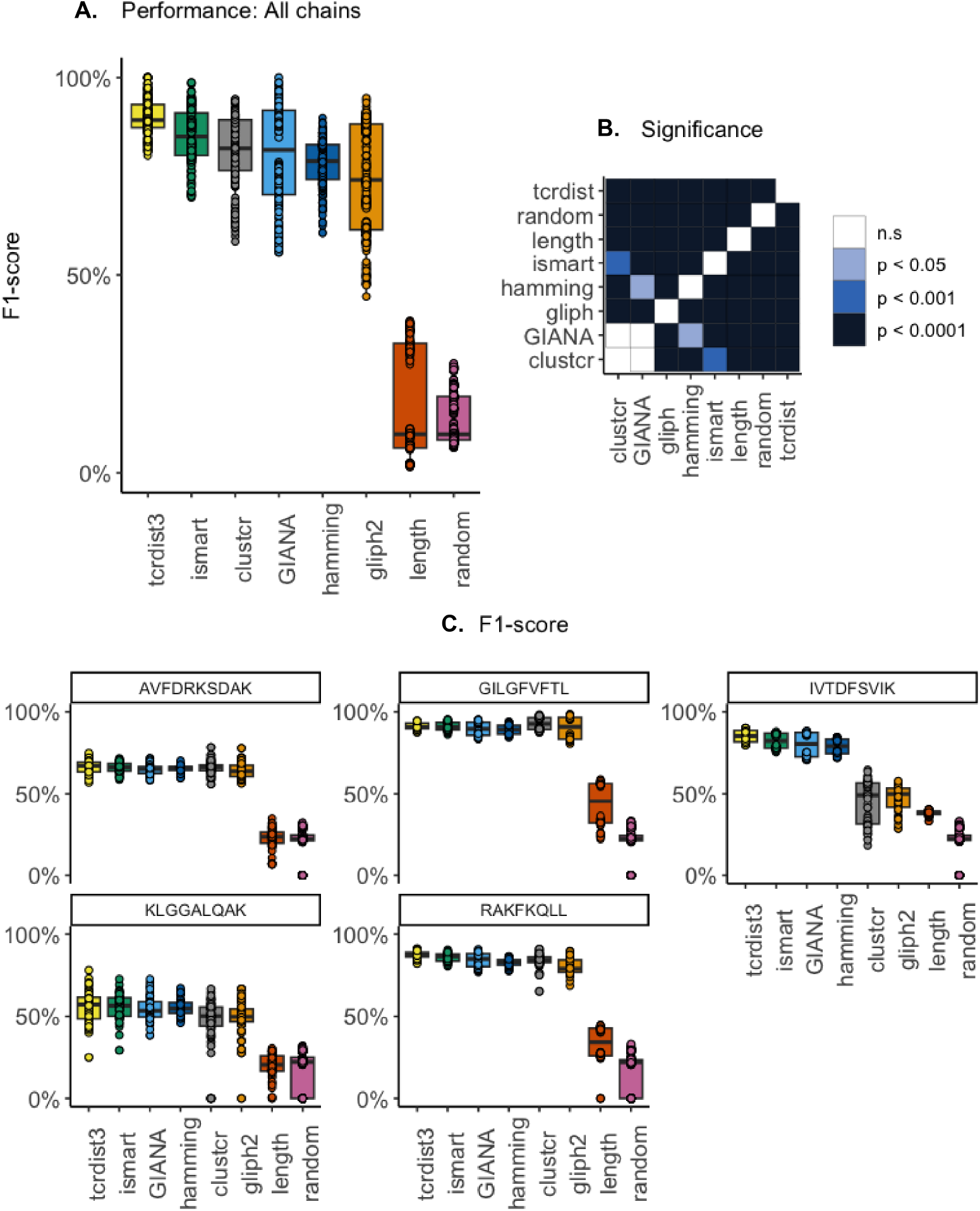
A-B) Comparative model performance for datasets V10-V1000, *α* and *β* chain selections combined. A) Predictive performance; B) Significance (p) values of comparisons between models following one-way ANOVA with *post hoc* Tukey’s HSD test; C) F1-score per epitope, dataset V500, *α* and *β* chain selections combined

An inspection of the size and purity of clusters generated by each model revealed high levels of similarity in the patterns produced by ClusTCR, GIANA, GLIPH2, iSMART, tcrdist3, and a Hamming model, which were in turn strikingly different from those produced by CDR3 length and random models (**Fig. 3A**). Notably, these six models produced large, pure clusters of thirty or more members in which the most frequent epitope was GILGFVFTL (Influenza A M-Protein) or RAKFKQLL (EBV BLZF1). *β* chain CDR3 motifs for the largest clusters associated with each of these epitopes were also near-identical except for length and random baseline models (**Fig. 3B**). Clusters in which AVFDRKSDAK (EBV EBNA-4), IVTDFSVIK (EBV EBNA-4), and KLQQALQAK (hCMV IE1) was the most common epitope were rarer, smaller, and less pure, producing less consistent CDR3 motifs (**Fig. S5**).

**Figure 3.**
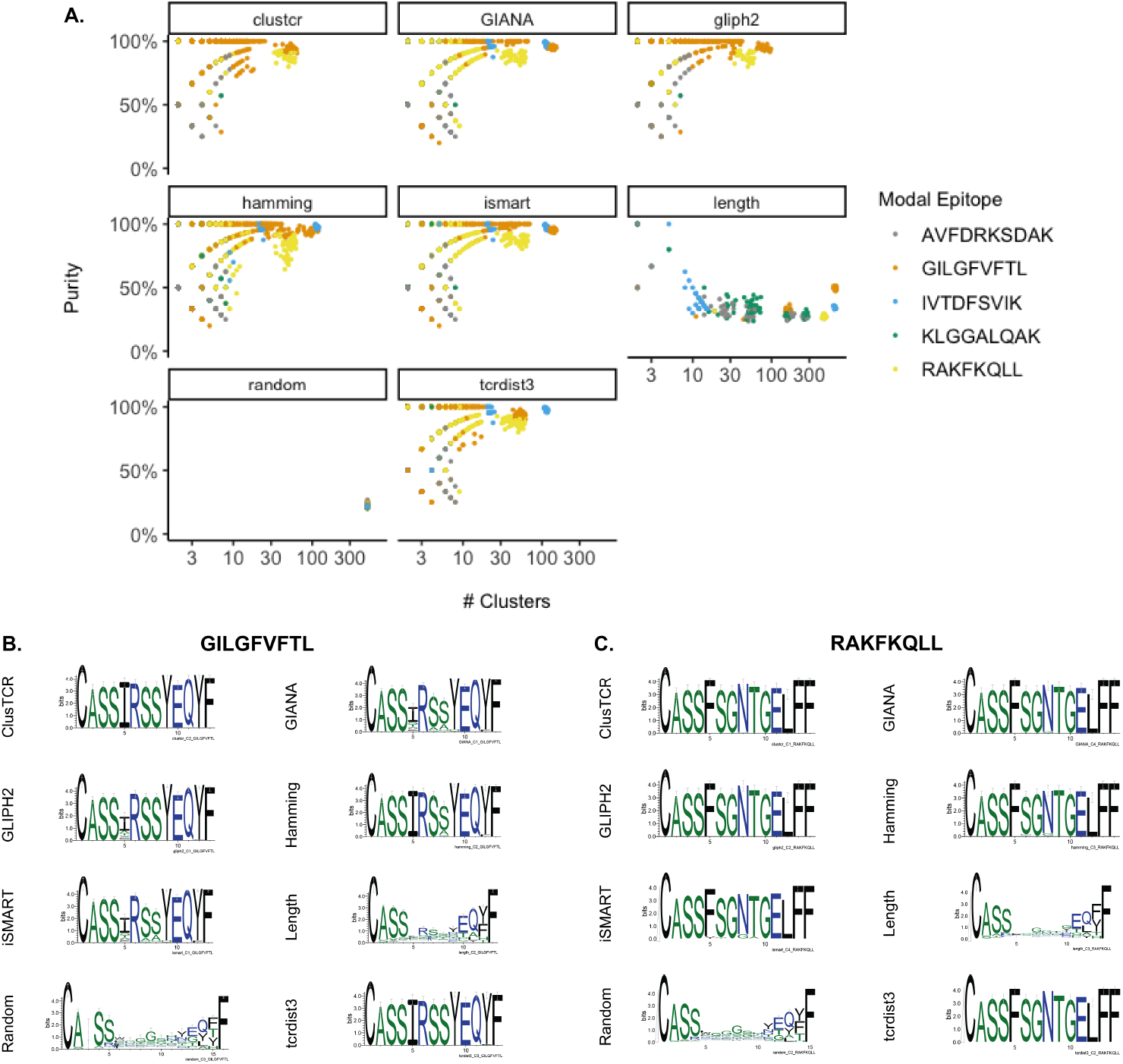
Qualitative analysis of UCM model outputs, for dataset V500 (*β* chain selection). A): Cluster purity and size distributions by modal epitope. B-C): Sequence logos for the largest clusters produced by a given model for a given epitope. B): GILGFVFTL; C): RAKFKQLL. Logos were produced with WebLogo [29] for TCRs in the largest cluster produced for a given model following sequence alignment with MUSCLE [30].

We next compared UCM computational speed by adding synthetic TCRs, produced with OLGA [31], to our input data (**Fig. 4A**). Here, the Hamming distance comparator outstripped the other models by a significant margin, slower only than length and random baselines. tcrdist3 scaled poorly: with memory constraints preventing use with repertoires greater than 10,000 TCRs for a single CPU and the majority of runtime being contributed by the computation of TCR distances (**Fig. S6**). Predictive performance was not materially impacted by the addition of 10,000 synthetic TCR sequences (**Fig. 4B** and **Table S6**). Taken together, these results suggest firstly that the variable performance gains observed for some models over a Hamming distance comparator come at the cost of a significant increase in runtime, at least when deployed over a single CPU. Secondly, all models were able to infer TCR epitope specificity materially better than a random comparator, even when labelled TCRs were diluted with synthetic TCRs at a ratio of 3:1.

**Figure 4.**
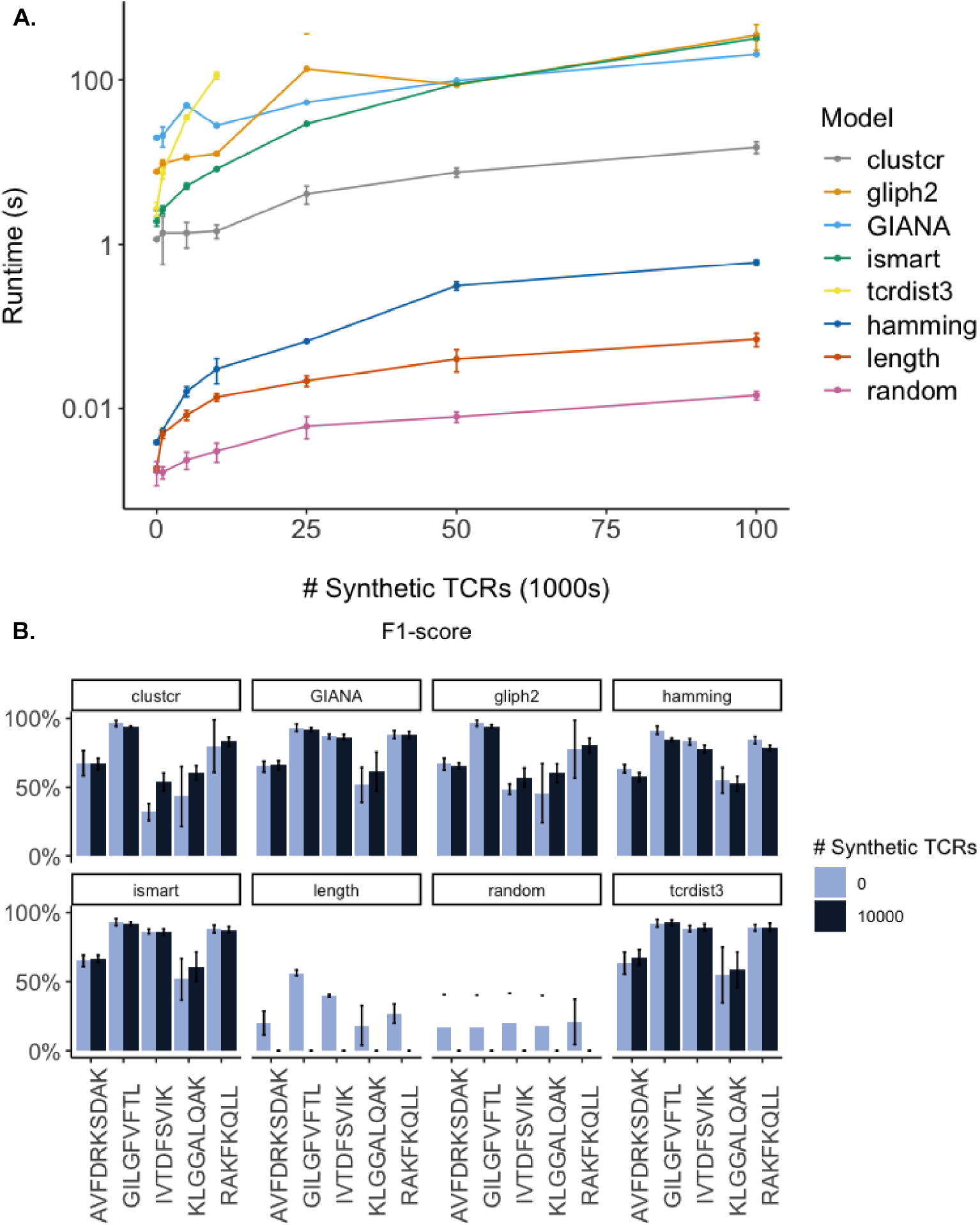
Investigating model scalability, comparing model runtimes as a function of the number of synthetic TCR sequences introduced with OLGA (Sethna et al., 2019). A) Runtimes. B): Epitope-specific F1-scores in the presence or absence of 10,000 synthetic TCR sequences. All experiments conducted on dataset V500 (Furthermore, rankings did not hold across *β* chain selection, 5 repeats).

## Discussion

Despite the exponential growth of orphan TCR datasets, and the widespread use of UCMs in de-orphanisation pipelines, an independent comparison off their predictive capacity has thus far been missing from the field. Here, we present a first, modest attempt to address this need. Our findings suggest that five commonly used UCMs show some variation in their ability to infer the specificity of a given set of TCRs, but that the best-performing models produce similar clusters to a simple Hamming distance model at considerably slower speeds. We first explore the possible factors underlying the observed performance differences, before reviewing the implications of these results for the challenging task of TCR specificity inference.

ClusTCR, GIANA, GLIPH2, iSMART and tcrdist3 consistently outperformed length and random baselines, however the relative rankings were sensitive to the chain selection, epitope, and dataset used (**Fig. 2, Fig. S3-4**). Intriguingly, global performance was generally within 1-3% of a simple Hamming distance model when accounting for variance (**Table S3**). One exception was tcrdist3, which achieved a 5% improvement in mean F1-score over the next best model when grouping analyses across datasets, epitopes and chain selections (**Table S3**). However, we cannot rule out that the observed performance gain for tcrdist is a product of the pre-processing and sampling strategy used, and we encourage further independent comparisons on other datasets. If real, one possible explanation of this improved performance is the use by tcrdist of inferred sequences for germline-encoded CDR1, CDR2, and pMHC-facing CDR2.5 regions of the TCR, instead of categorical representations of the corresponding gene code [14]. Indeed, a simple predictive model combining tcrdist and a K-nearest neighbour model achieved superior performance to many DNN-SPMs using gene codes in a recent benchmarking exercise [12].

Although recent structural [32], statistical [33], and predictive [9] analyses suggest that both polypeptide chains play an important role in epitope recognition, we observed consistently lower F1-scores for *α* compared to *β* chain selections (**Table S3**). One possible explanation is that the default model hyperparameters have been optimised for *β* chain data, which make up the majority of published TCR-epitope pairs [8].

Alternatively, the *β* chain may simply contribute more to determination of overall epitope specificity, as a product of its increased diversity relative to *α* chains, however this warrants further investigation. For example, modelling strategies that permit integration of *α* and *β* chain pairing with transcriptomic and phenotypic information, including graph network approaches such as CoNGA [34], may help efforts to decode the relative contribution of chain pairing to epitope specificity at single cell resolution.

If model performance is sensitive to the choice of dataset and pre-processing strategy, and the five UCMs produce similar clusters to a Hamming distance model, how then can one decide which UCM to use for analysis and/or co-clustering of large TCR sequence datasets? One important lens is scalability, but here again a Hamming distance model performed best, apart from length and random baselines (**Fig. 4A**). Another consideration is accessibility. We note for example that at present only GLIPH2 is currently available as both a web tool and command line executable, the other models requiring some familiarity with programming languages such as Python and R.

Finally, what do these results tell us about the relative capacity of UCMs for inference of orphan TCR specificity? Combining labelled instances and synthetic sequences produced with OLGA (**Fig 4B** and **Table S5**) provides a window into model performance in co-clustering of reference and orphan TCRs. Encouragingly, we see that UCMs are able to successfully cluster TCRs of common specificity in the presence of a synthetic background of 10,000 TCRs. Our results therefore support the continued use of these models in de-orphanisation pipelines, by co-clustering labelled and unlabelled TCRs, an approach that could theoretically be applied to both seen and unseen epitopes.

## Study Limitations

The significant scientific and economic potential of a generalisable solution to prediction of TCR epitope specificity has encouraged the development of a multitude of new SPMs and UCMs, summarised in [6]. However, the scope of the present study is limited to a handful of commonly used UCMs, on the basis of their widespread use and relative freedom from training data bias as compared to DNN-UCMs and SPMs. Nonetheless, an independent comparison of UCMs, DNN-UCMs and DNN-SPMs would be of great use to the community. There is also growing evidence that inclusion of both *α* and *β* chains improves predictive performance in SPMs and DNN-SPMs. However, whilst ClusTCR, tcrdist3, and GLIPH2 may all theoretically be applied to paired chain data, cluster assignments are produced for a given CDR3 independent of the other chain for all but tcrdist3. An investigation of whether the integration of *α* and *β* chain information improves performance equally across models might reveal the relative merits of each, when applied to large scale single-cell experiments. Time and technical limitations prevented extension of the present analysis to comparative performance over parallel CPUs, and to the GPU-enabled versions of ClusTCR and GIANA. Finally, whilst we have made efforts to investigate relative model performance under a variety of pre-processing conditions, predictive power is sensitive to the dataset and pre-processing methodology. Therefore, an extension to other public datasets, as well as to complete repertoire data from large population studies such as [35], would add certainty to the conclusions drawn.

## Acknowledgments

Our comparative framework used borrowed heavily from that developed by the Meysman group for ClusTCR (https://github.com/svalkiers/clusTCR), to whom the authors express their gratitude. Thanks are also due to Dr Ricardo A. Fernandes and Sam Farrar for critical review. H.K. is supported by funding from the UK Medical Research Council grant number MC_UU_12010/3. D.H. receives administrative and financial support from the Biotechnology and Biological Sciences Research Council (BBSRC) (grant number BB_T008784_1) and from the Rosalind Franklin Institute.

## Ethics statement

D.H. provides consultancy services to companies active in T cell antigen discovery and vaccine development. The other authors declare no competing interests.

## Methods

### Datasets

A consolidated dataset of paired TCR amino acid sequences of human origin was developed using instances drawn from VDJdb [24] and from McPas-TCR [36] as a separate test set. Sequences derived from a 10X study of healthy donors, and CDR3 sequences containing non amino acid symbols, were removed from the input data. V and J gene codes were processed for consistency with IMGT reference sequences [37].

Duplicates were removed within and between datasets using CDR3-V-J bio-identities for both *α* and *β* chains, such that a given TCR was encoded in the format CDR3 *α* TRAV TRAJ CDR3*β* TRBV TRBJ. Only those TCRs having TRA or TRB genes included in the reference IMGT alleles of the tcrdist module of the CoNGA conda package (v.0.1.1) were retained, to ensure that consistent numbers of sequences were provided to each model. Benchmarking experiments were performed on VDJdb data after selection and down sampling as described in **Results**. 343

### Models

A systematic review of the literature was conducted to identify studies presenting novel methods for prediction of antigen specificity from TCR sequences. ClusTCR [25], GIANA [26], GLIPH2 [17], iSMART [20], and tcrdist3 [27] were shortlisted for analysis based on the availability of open-source python packages or executable files. ALICE [38] was excluded as more appropriately applied to the identification of expanded clones in individual patient repertoire data. Background methodological detail is included for each of the selected algorithms in **Section S1**. The analytical framework developed to accompany the ClusTCR package was adapted to permit comparison of each of the models described below. All benchmarking experiments were run on a single remote Intel(R) Xeon(R) CPU (E7-8891 v3 @ 2.80GHz) to ensure fair comparison of algorithms with and without parallel processing capability.

The ClusTCR python package (v1.0.2) was imported with Anaconda and implemented using default settings. Benchmarking of ClusTCR was conducted with the CPU version for fair comparison with non-parallelisable models. GLIPH2 was downloaded from the developers’ website and run using a combined CD4/CD8 reference, otherwise using default parameters. Where a given sequence was assigned to more than one putative cluster, absolute cluster assignments were made to the cluster having the greatest probability in the output. A Hamming distance model was adapted from a version published in the ClusTCR repository which makes use of sequence hashing for efficient CDR3 comparison, first grouping CDR3 sequences by length and then sorting these superclusters into subclusters with a Hamming distance of 1. iSMART was implemented as in [25] except that V gene usage was included by default. GIANAv4.1 was downloaded from GitHub with an IMGT TRBV reference and implemented in CPU mode using default settings following the framework developed for iSMART. tcrdist3 (v0.2.2) was installed with PyPI and called with a Python script making use of sparse distance matrices for large datasets. tcrdist3 amino acid distance matrices were generated with the default meta-clonotype radius of 50 and clustered with DBScan (eps=0.5) after an initial parameter search (**Fig. S1**). A faster C++ implementation of tcrdist is available as part of the CoNGA package [34], however a steep drop-off in epitope-specific performance was observed when combining this model with DBSCAN (**Fig. S1**). Greedy clustering, used in the original tcrdist publication [14] and evaluated in [25], was excluded from the analysis due to prohibitively slow runtimes. Finally, length and random baseline models were added which assigned TCRs to clusters based on CDR3 amino acid sequence length and random shuffling, respectively.

### Metrics

Performance of each model was analysed using the cluster metrics described previously in [25], [39]: purity, consistency and retention, with the addition of the scikit learn implmentation of adjusted mutual information (AMI)[28] to account for cluster entropy.

Balanced accuracy, weighted precision, weighted recall, and weighted F1-score were computed as a mean over all clusters and for a given epitope, using the Scikit learn library [28].

### Statistics

All statistical comparisons were performed using an R implementation of one-way analysis of variance (ANOVA) and Tukey’s HSD for post-hoc significance testing, which analyses are accessible in the accompanying GitHub repository. Comparative boxplots and probability heat maps were produced using ggplot2[40].

### CDR3 amino acid motifs

Sequence logos were produced from *β* chain selections of dataset V500 by retaining TCRs having the modal length from the largest cluster for each of five epitopes of interest. Sequences were aligned with MUSCLEv5.1 [30], and logos produced from the resulting multiple sequence alignments with WebLogo v3.7.12 [29].

## Supplemental Information

### Section S1: Model methodologies

Here we provide a brief overview of the principal methods underlying each of the models tested, referring the interested reader to the original citations for further details.

**ClusTCR** [25] makes use of a two-step approach to clustering, in which an N x M matrix of CDR3 amino acid sequence and physicochemical properties is sorted into superclusters using the Faiss library, and the resulting embeddings are sorted with KMeans. A graph network of distances is then produced from these superclusters based on Hamming distances between length sorted CDR3 sequences. Final cluster assignments are made by applying Markov Clustering (MCL) to the network graph.

**GIANA** [26] applies multidimensional scaling (MDS) to produce matrix representations of TCR CDR3 sequences that approximate BLOSUM62 physicochemical properties, such that the Euclidean distance between two sequences represented with MDS is equivalent to the Smith-Waterman alignment between the BLOSUM representations of those sequences. MDS vectors are pre-sorted on length, and the resulting superclusters are then sorted into subclusters using the Faiss library before clustering on Smith-Waterman distances between kmers.

**GLIPH2** [17] is an update to GLIPH [15] that combines global and local cluster analyses. Global distance is defined as sequence mismatches in CDR3 sequences differing at a given position according to a BLOSUM62 subsititution matrix, having shared TRBV gene usage and identical length. Local distance is computed as a statistically significant kmer frequency enrichment in residues predicted to contact peptide-MHC, compared to a sample population. **iSMART** [20] incorporates CDR3 and (optionally) V gene usage information, pre-sorting CDR3 sequences according to length and imposing a gap penalty for length mismatched CDR3s related by a single insertion. Alignment scores are computed for a subset of the CDR3 sequences using a BLOSUM62 substitution matrix, and output clusters are assigned based on a threshold alignment score.

**tcrdist3** [27] is the latest iteration of tcrdist [14], which makes use of a BLOSUM62 mismatch distance between CDR1, CDR2, CDR2.5 (an MHC-facing loop), and CDR3 sequences. Non CDR3 sequences are inferred from a reference database, a gap penalty is applied to account for sequence insertions/deletions, and a combined similarity score is computed that assigns greater weighting to CDR3 sequences. The resulting distance matrix may then be clustered, for example using a greedy hierarchical search (see **Methods** and **Fig. S1**).

## Supplemental Tables

**Table S1.** Dataset size.

| Dataset | Minimum TCRs per epitope | # Unique epitopes | N total |
| --- | --- | --- | --- |
| *VDJdb* |  |  |  |
| V10 | 10 | 76 | 760 |
| V50 | 50 | 25 | 1250 |
| V100 | 100 | 16 | 1600 |
| V500 | 500 | 5 | 2500 |
| V1000 | 1000 | 4 | 4000 |
| *McPas-TCR* | N/A | 24 | 509 |

**Table S2.** Frequency of TCR representatives per epitope in preprocessed VDJdb input data prior to down sampling.

| Epitope | # Instances |
| --- | --- |
| KLGGALQAK | 13,552 |
| GILGFVFTL | 1,830 |
| AVFDRKSDAK | 1,143 |
| RAKFKQLL | 1,120 |
| IVTDFSVIK | 572 |

**Table S3.**
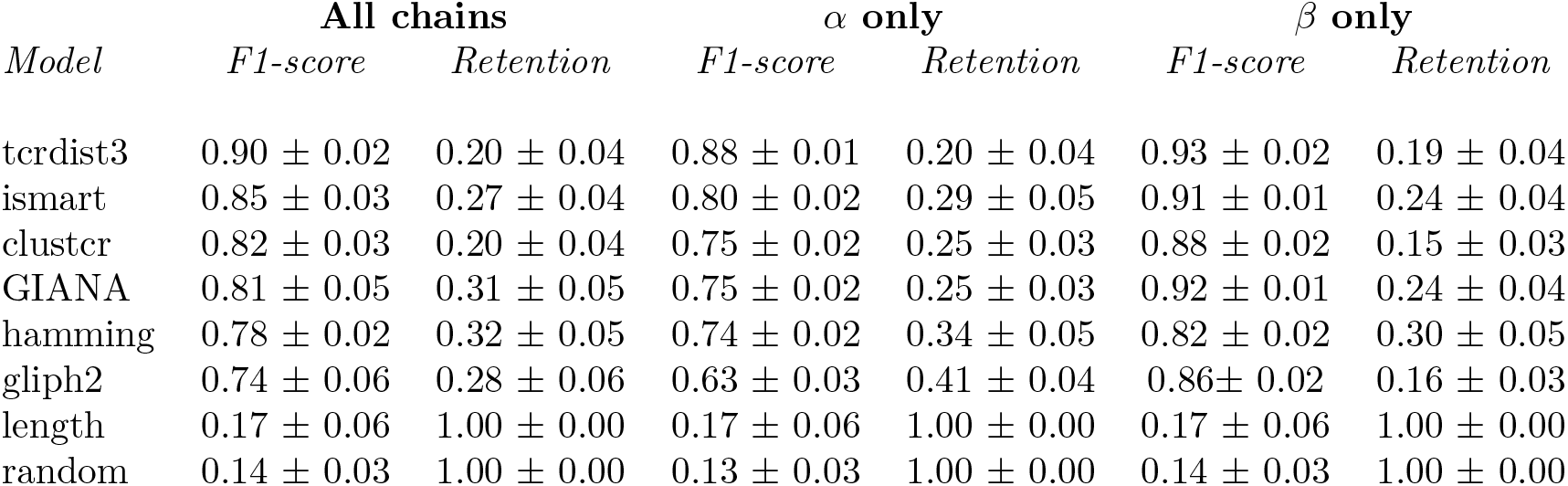
Global UCM performance, showing mean values ± 95% confidence (datasets V10, V50, V100, V500 and V1000 combined, *α* and *β* chain selections, 25 repeats.)

**Table S4.**
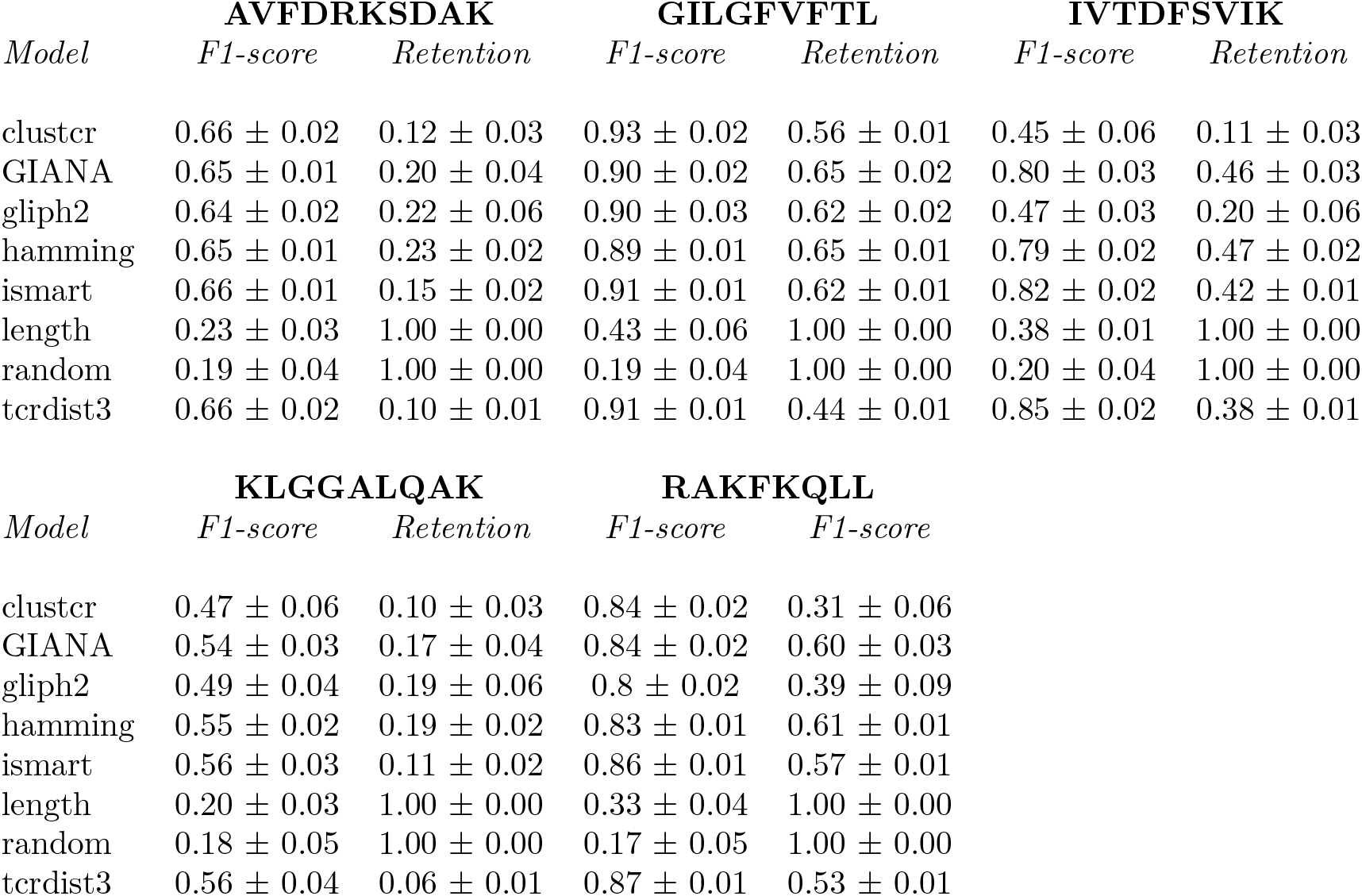
UCM performance and retention by epitope, showing mean values ± 95% confidence (dataset V500, *α* and *β* chain selections, 25 repeats.)

**Table S5.**
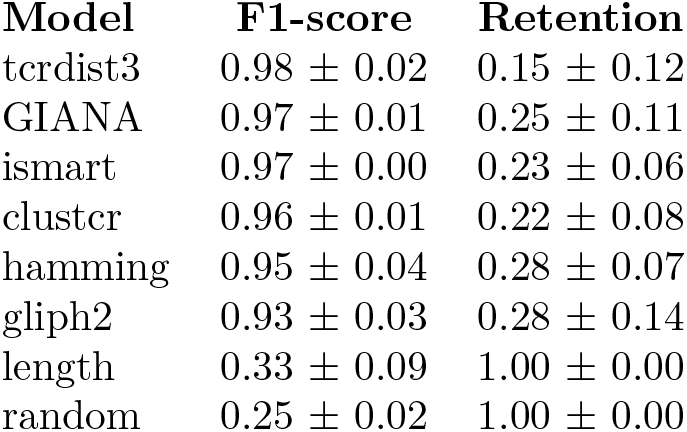
UCM performance on instances from McPas-TCR (5 repeats).

**Table S6.**
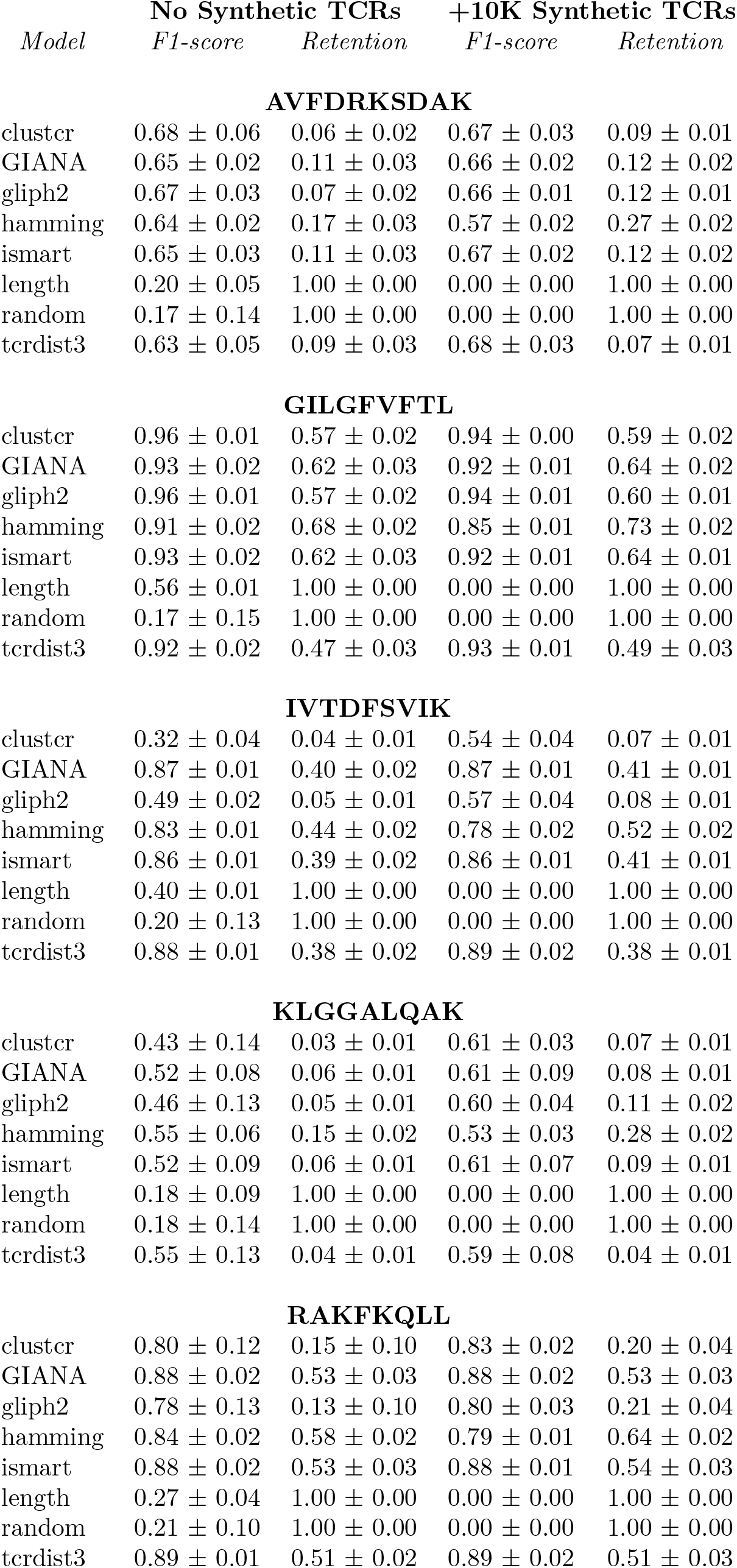
UCM performance in the presence of synthetic TCR sequences produced with OLGA, V500, *β* chain selections (5 repeats).

## Supplemental Figures

**Figure S1.**
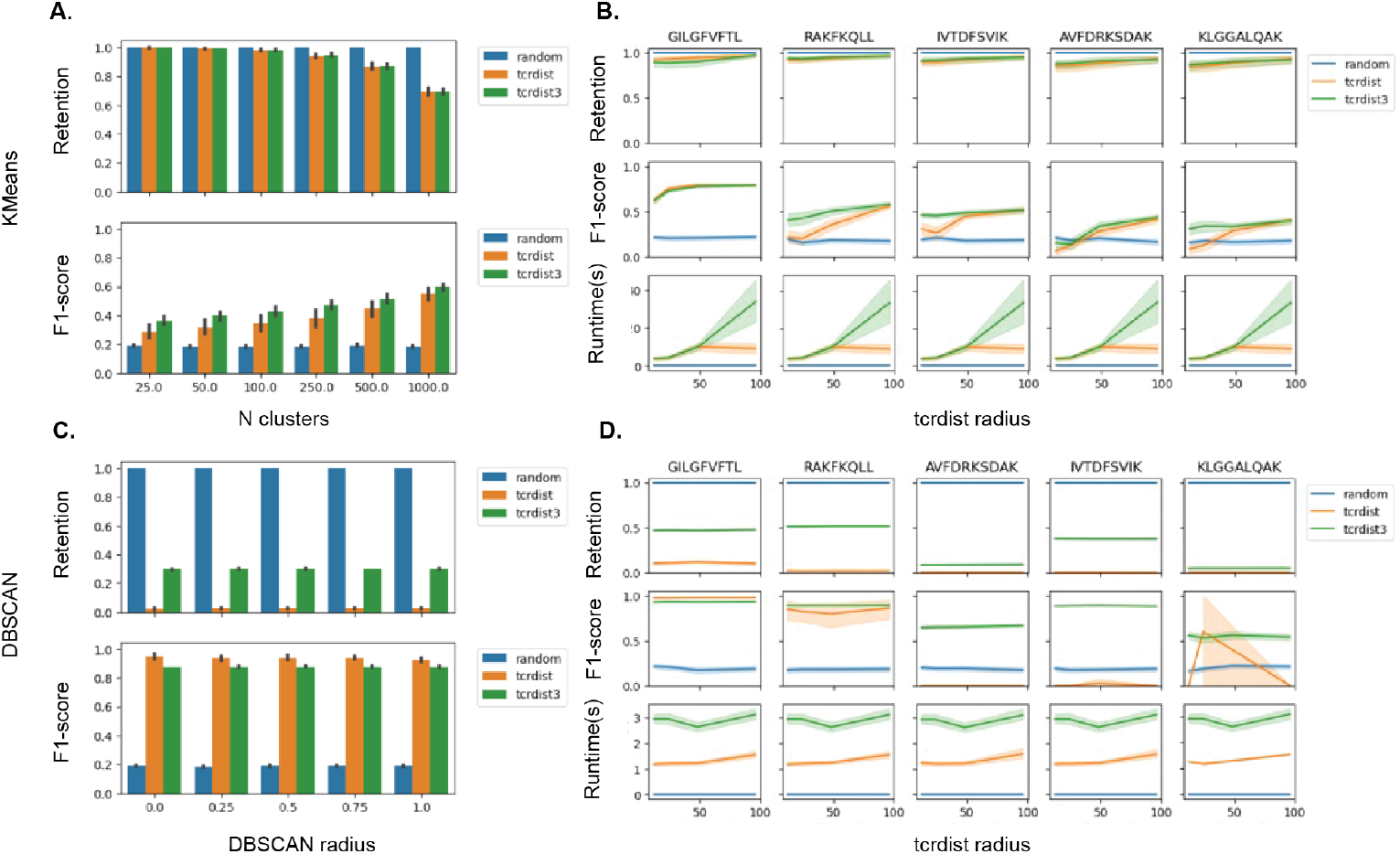
Selecting tcrdist hyperparameters for C++ (tcrdist) or python (tcrdist3) implementations of tcrdist (Schattgen et al., 2022, Mayer-Blackwell et al., 2021), using KMeans (A-B) or DBSCAN (C-D) applied to dataset V500, *β* chain selections. A), C): Performance as a function of the number of clustering algorithm hyperparameters. B, D): performance per epitope as a function of tcrdist radius.

**Figure S2.**
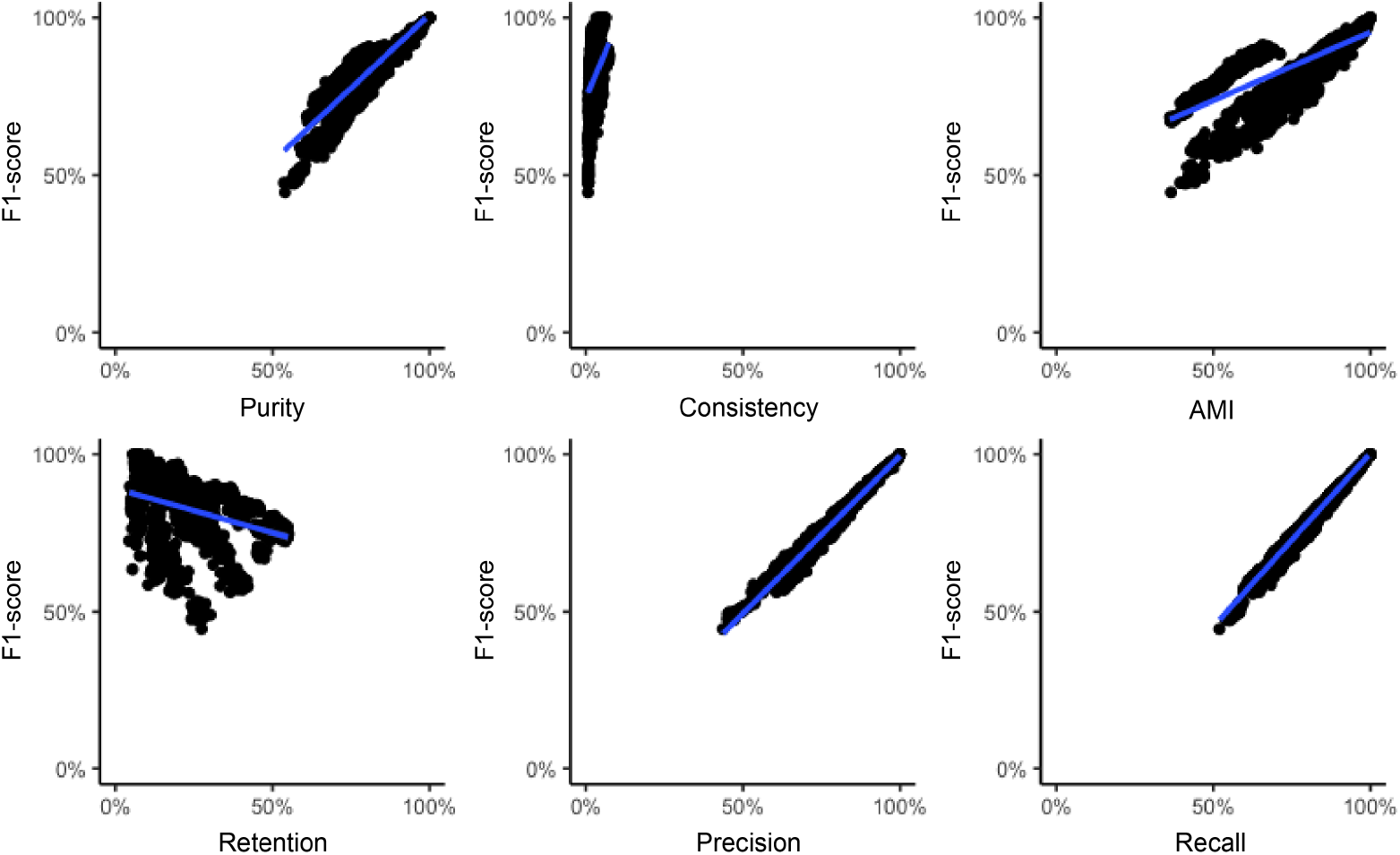
Correlation of UCM metrics, datasets V10, V50, V100, V500 and V1000, *α* and *β* chain selections combined (25 repeats).

**Figure S3.**
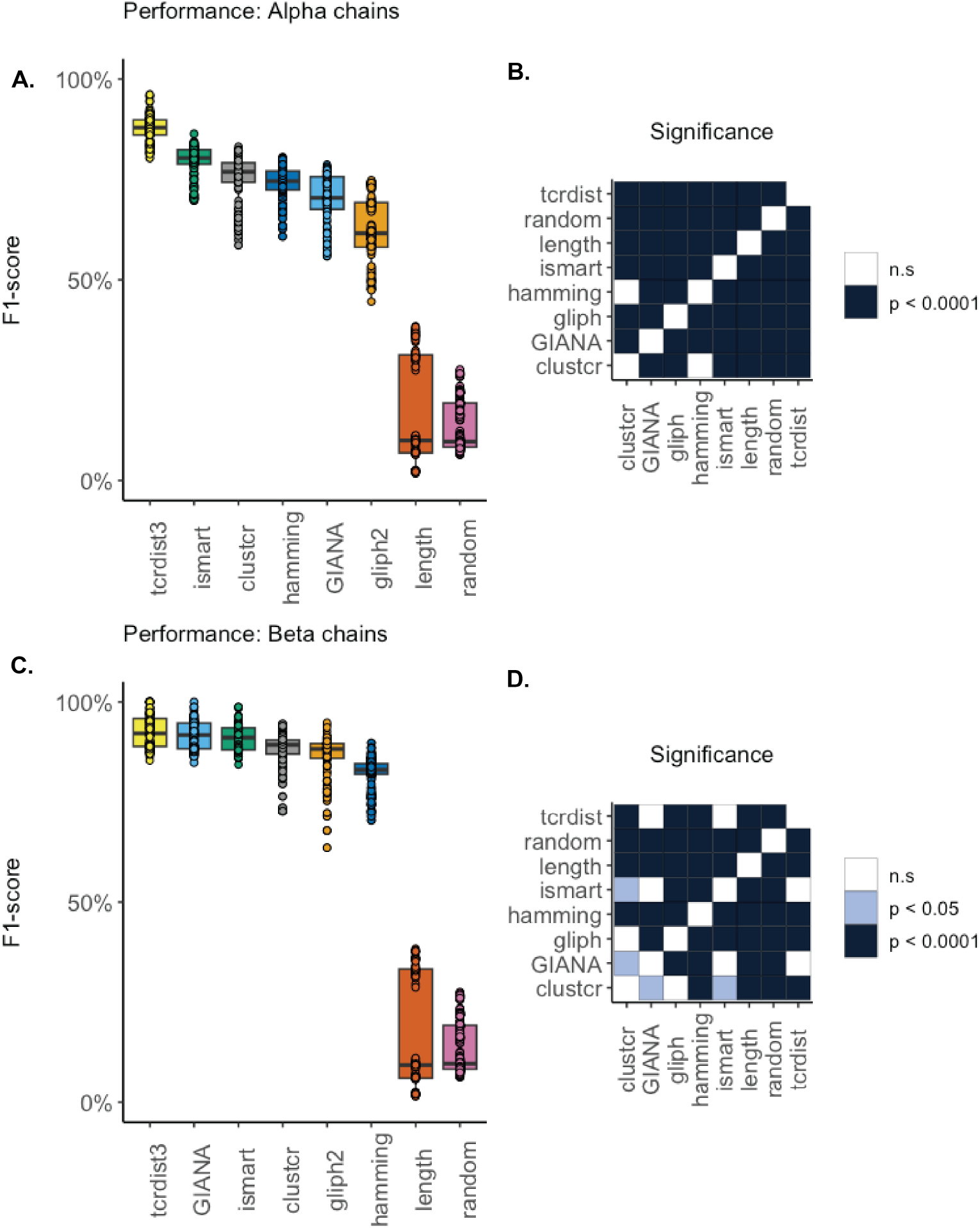
Performance differences (A, C) and statistical significance (B, D) by chain selection, datasets V10, V50, V100, V500 and V1000 combined (25 repeats). A-B): *α* chain selections; C-D) *β* chain selections.

**Figure S4.**
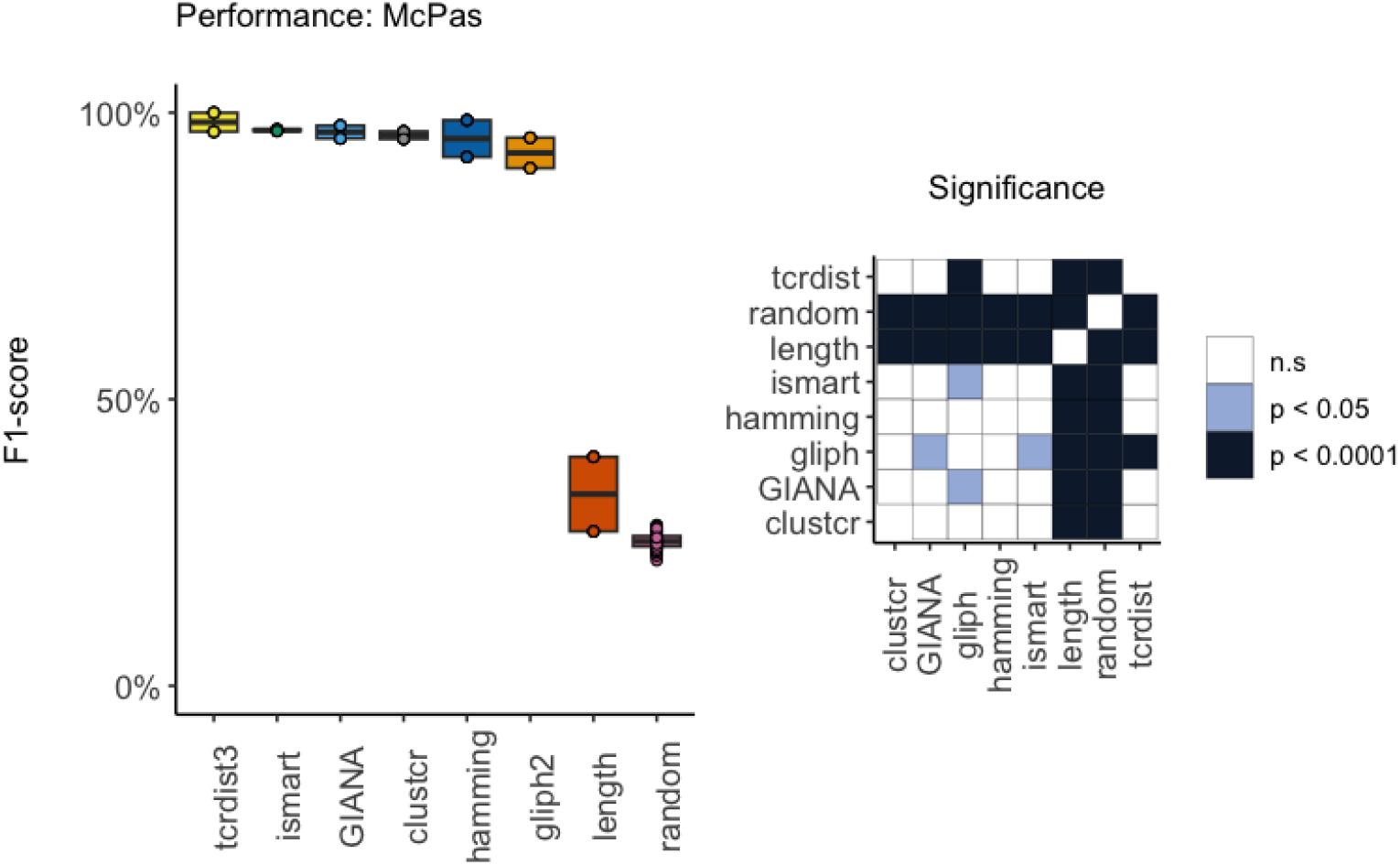
Performance differences (*left*) and significance (*right*) for McPas-TCR, *α* and *β* chain selections (25 repeats).

**Figure S5.**
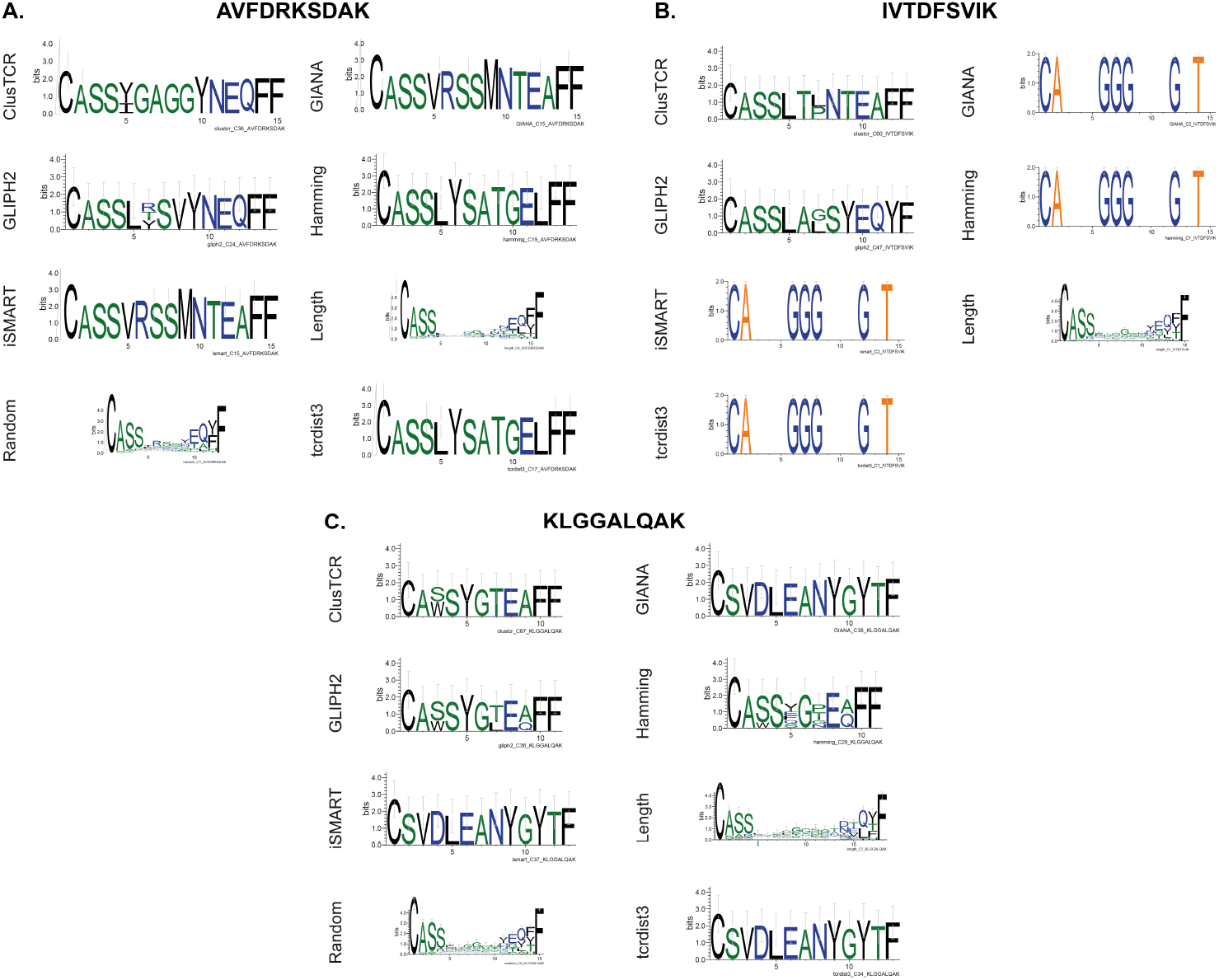
Motifs for the largest cluster formed per epitope, dataset V500 (*β* chain selection). A): AVFDRKSDAK; B): IVTDFSVIK; C): KLGGALQAK

**Figure S6.**
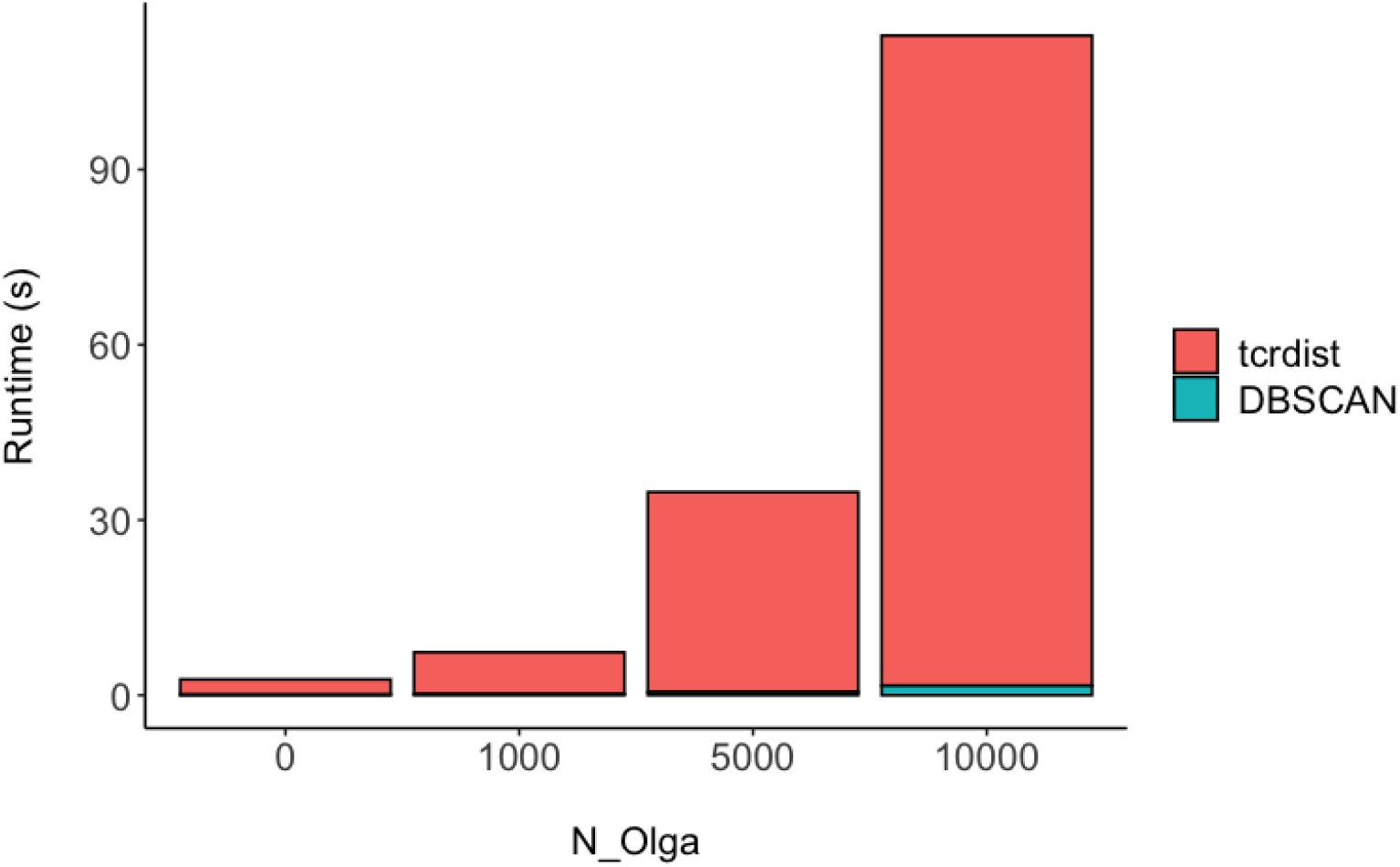
Relative contribution to runtimes of tcrdist3 matrix calculation and clustering with DBSCAN in the presence of increasing synthetic TCR sequences produced with OLGA[31], dataset V500, *β* chain selection [31].

